# Reducing stomatal density by expression of a synthetic EPF increases leaf intrinsic water use efficiency and reduces plant water use in a C_4_ crop

**DOI:** 10.1101/2024.02.01.578512

**Authors:** John N. Ferguson, Peter Schmuker, Anna Dmitrieva, Truyen Quach, Tieling Zhang, Zhengxiang Ge, Natalya Nersesian, Shirley J Sato, Tom E. Clemente, Andrew D.B. Leakey

**Affiliations:** Institute for Genomic Biology; Department of Plant Biology; Department of Crop Sciences, University of Illinois at Urbana-Champaign, Urbana, Illinois; Depatment of Agronomy and Horticulture, University of Nebraska, Lincoln, Nebraska; School of Life Sciences, University of Essex, Colchester, UK

**Keywords:** *Sorghum bicolor*, stomata, C_4_ photosynthesis, water-use efficiency

## Abstract

Enhancing crop water use efficiency (WUE) is a key target trait for climatic resilience and expanding cultivation on marginal lands. Reducing stomatal conductance (*g_s_*) through manipulating stomatal density has been observed to translate to improved WUE in multiple C_3_ crop species. However, reducing *g_s_* in C_3_ species often reduces photosynthetic carbon gain. A different response is expected in C_4_ plants because they possess specialized anatomy and biochemistry which concentrates CO_2_ at the site of fixation. This modifies the photosynthesis (*A_N_*) relationship with intracellular CO_2_ concentration (*c_i_*) so that photosynthesis is CO_2_-saturated and reductions in *g_s_* are unlikely to impair *A_N_*. To test this hypothesis, genetic strategies were investigated to reduce stomatal density in the C_4_ crop sorghum. Constitutive expression of a synthetic epidermal patterning factor (EPF) transgenic allele in sorghum, lead to reduced stomatal densities. A moderate reduction in stomatal density did not strengthen stomatal limitation to A_N_, improved WUE, reduced water use, and avoided loss of carbon fixation during a period of water deprivation. However, these positive outcomes were associated with negative pleiotropic effects on reproductive development and photosynthetic capacity. Avoiding pleiotropy by targeting expression of the transgene to specific tissues provides a potential pathway to optimal agronomic outcomes.

## Introduction

Water availability is a critical factor limiting crop productivity worldwide (Boyer 1982). Water Use Efficiency (WUE) has been recognized as an important trait for crop improvement for over a century (Briggs and Shantz 1917) because it describes how much productivity can be achieved per unit of water used by the crop. WUE is key to terrestrial plant function because it reflects the inevitable trade-off between losing water vapor from leaves to the atmosphere, while stomata are open, to allow photosynthetic CO_2_ uptake. Climate change is impacting precipitation patterns and vapour pressure deficit, which in turn impact water availability and demand in cropping systems (Sadok *et al*. 2021, Chiang *et al*. 2021). *In silico* modelling that incorporates historical yields and projected environmental conditions suggests that drought events will continue to be a key driver of yield losses in the future (Webber *et al*. 2018). Water supplies for irrigation are limited and unsustainable (WWAP 2015). Demand is increasing for agricultural products from marginal lands (Gelfand *et al*. 2013; Khanna *et al*. 2021). Consequently, there is renewed focus on whether crop productivity, sustainability and resilience can be enhanced through improvements to WUE (Hatfield and Dold, 2019; Leakey *et al*. 2019; Bailey-Serres *et al*. 2019; Sales *et al*. 2021).

Intrinsic WUE (iWUE) is defined as the ratio of the rate of net photosynthetic CO_2_ assimilation (A_N_) relative to stomatal conductance (*g_s_*). Improving iWUE can be achieved by increasing A_N_ without a matching increase in *g_s_*, or by decreasing *g_s_* without a matching decrease in A_N_. However, many studies of natural and engineered genetic variation within a broad diversity of plant species have shown A_N_ and *g_s_* to be correlated (Leakey *et al*. 2019; Deans *et al*. 2020). Consistent with this expectation, in C_3_ species, theory and experimentation have shown that a decrease in stomatal density drives lower *g_s_* and improves iWUE, but often at the cost of lower A_N_ (Wang *et al*. 2016; Hughes *et al*. 2017; Mohammed *et al*. 2019; Caine *et al*. 2019; Dunn *et al*. 2019). So, de-coupling of A_N_ and *g_s_* in transgenic plants is rare and, when it is achieved, iWUE is often reduced rather than increased (Flexas *et al*. 2013).

Meanwhile, crop genotypes selected by breeders for greater iWUE have often been found to be innately less productive, as part of a generally conservative growth syndrome (Condon *et al*. 2004). Although successes have resulted from screening stable carbon isotopes as a proxy to select for high iWUE in C_3_ species (Rebetzke *et al*. 2006), scepticism remains about the potential for meaningful improvement of WUE without a concomitant hit on yield (Condon *et al*. 2004; Blum 2005; Sinclair 2012).

Recent innovations have emerged that open the door for improving WUE and productivity in C_4_ crops through lower *g_s_* via reduction in stomatal density. These include novel high-throughput phenotyping tools to capture stomatal density and WUE traits that are accelerating the speed and scale of experimentation (Ferguson *et al*. 2021, Pignon *et al*. 2021, Xie *et al*. 2021). Importantly, unlike C_3_ crops, C_4_ feedstocks possess a carbon concentrating mechanism that increases [CO_2_] in the bundle sheath cells where the primary photosynthetic enzyme Ribulose-1,5-Bisphosphate Carboxylase Oxygenase (RuBisCO) is located; resulting in concentrations significantly greater than atmospheric [CO_2_] (von Caemmerer and Furbank 2003). Consequently, the relationship between A_N_ and intercellular [CO_2_] (c_i_) features a much steeper initial slope and a sharper inflection point than that observed in C_3_ species (Leakey *et al*. 2009). Atmospheric [CO_2_] has risen from 370 ppm in 2000 to 417 ppm in 2023 (https://gml.noaa.gov/ccgg/trends/). As a result, the c_i_ at which photosynthesis operates in C_4_ crops is very close to saturation, which raises it above the A/ci curve inflection point (Leakey *et al*. 2006; Ghannoum, 2008; Markelz *et al*. 2011). This means more CO_2_ is entering the leaf than is required to maintain A_N_ and, theoretically, *g_s_* could be reduced to increase iWUE with little (<2%) to no increase in the stomatal limitation of A_N_ (Leakey *et al*. 2019). The magnitude of reductions in stomatal density needed to achieve an optimal *g_s_*is unknown, but too large a reduction in stomatal density would lower ci to the point that it would fall below the inflexion point on the A/ci curve, where stomatal limitation to A_N_ becomes substantial (Leakey *et al*. 2019). Mechanistic crop modelling suggests that a genetic strategy that translates to a reduction in *g_s_* by 20% would significantly increase yields of a C_4_ feedstock, like sorghum, with the greatest gains occurring in marginal land locations where yields are currently low (Leakey *et al*. 2019). Overexpression of STOMATAL DENSITY AND DISTRIBUTION 1 (SDD1) in maize provided evidence in support of this approach, driving reduced stomatal density and g_s_, without a decrease in A_N_ (Liu *et al*. 2015). But, pleiotropy was observed, with the low stomatal density plants also having greater photosynthetic capacity than WT. The consequences of over-expressing SDD1 for development, productivity, and WUE at the whole-plant scale were not reported.

Whole-plant transpiration rates regulate passive nitrogen uptake from the soil (Niu *et al*. 2007; Matsunami *et al*. 2010; Kunrath *et al*. 2020). Therefore, a potential drawback of a genetic strategy to improve WUE via a reduction in *g_s_* is that nitrogen flux in the transpiration stream might be reduced, which in turn would negatively impact photosynthetic capacity and productivity. A significant fraction of leaf nitrogen is invested in photosynthetic proteins, with C_4_ species allocating less nitrogen to Rubisco than C_3_ species, but more nitrogen to other soluble proteins and thylakoid components (Ghannoum *et al*. 2011). However, with ectopic expression of EPF1 in the C_3_ species wheat, barley and rice, photosynthetic capacity was reported not to decline (Hughes *et al*. 2017; Caine *et al*. 2019; Dunn *et al*. 2019).

Testing the proposed approach to developing C_4_ crops with greater WUE is aided by the discovery of a network of genes that regulate leaf epidermal cell fate and, thereby, stomatal density in C_3_ species, because their C_4_ orthologs can be used as initial candidate genes for testing (McKown and Bergmann 2020). Most studies that have manipulated stomatal density in C_3_ species have achieved this by overexpressing the native form of EPIDERMAL PATTERNING FACTOR 1 (EPF1), a negative regulator of stomatal development (Harrison *et al*. 2020), or down-regulating EPIDERMAL PATTERNING FACTOR-LIKE 9 (EPFL9 or stomagen), a positive regulator of stomatal development (Karavolias *et al*. 2023). The EPF family of secreted signaling peptides function within the epidermal cell layer to regulate stomatal patterning (Hara *et al*. 2007; Hunt and Gray, 2009). Various EPFs fusions swapping the loop and scaffold regions of EPF2 and EPFL9 in Arabidopsis demonstrated that the loop region confers the functional specificity of EPFs (Ohki *et al*. 2011). EPF peptides trigger a mitogen activated protein (MAP) kinase signaling pathway that regulates the stability of SPEECHLESS (SPCH). SPCH is a basic helix-loop-helix (bHLH) transcription factor that contributes to the determination of cell division and fate transitions (Lau *et al*. 2014). MAP kinases phosphorylate and destabilise SPCH preventing the initiation of stomatal lineage cells, consequently overexpressing EPF genes negatively regulates stomatal density via a decrease in SPCH protein levels (Kumari *et al*. 2014). Recent evidence from C_3_ monocotyledonous species, such as barley, rice, and wheat, suggests that despite substantial differences in leaf expansion and stomatal development, native grass EPFs act in a similar way to the dicotyledonous model Arabidopsis’ EPFs; regulating entry to and progression through the stomatal cell lineage (Hughes *et al*. 2017; Mohammed *et al*. 2019; Caine *et al*. 2019; Dunn *et al*. 2019). On the other hand, the role of other regulators of stomatal and leaf development are not always strictly conserved across species and few stomatal development genes have been tested in C_4_ species (Liu *et al*. 2009; Raissig *et al*. 2016; Rassig *et al*. 2017; Schuler *et al*. 2018; Hughes and Langdale 2022).

Sorghum has high WUE and is well adapted to xeric environments (Rooney, 2014). It is an important model C_4_ species, with significant contributions to the bioeconomy as a feedstock for food, feed and industrial applications (Paterson *et al*. 2009; Morris *et al*. 2013; Castro *et al*. 2015). Various models have shown that in high-yielding environments, a reduced water use trait can protect yield during soil moisture deficits (Sinclair *et al*. 2005; Truong *et al*. 2017).

This study tested two genetic designs to reduce stomatal density in sorghum, in order to address the knowledge gap about how engineering reduced stomatal density would impact photosynthetic physiology and whole-plant function in a C4 species. In the first design, the native sorghum EPF1 (SbEPF1) was constitutively expressed under control of the sugarcane ubiquitin4 (Ubi4) promoter (Wei *et al*., 2003). In the second design, a fusion element was synthesized that combined elements of the sorghum orthologs of AtEPF2 and AtEPFL9 (SbEPF_syn_) and placed under control of the Ubi4 promoter. Comparing sorghum events with reduced stomatal density verses wildtype controls addressed whether reduced stomatal density: i) can drive reductions in *g_s_* without significantly increasing stomatal limitation to A_N;_ ii) drives pleiotropic effects on leaf physiology or plant development; iii) produces leaf-level reductions in *g_s_* that scale to reduced whole-plant water use; and iv) alters whole-plant biomass production.

## Materials and methods

### Assembly of binary vectors

The leucine rich repeat, receptor like kinase ERECTA corresponding interacting partner EPIDERMAL PATTERNING FACTOR (EPF), EPF1, is a negative regulator of stomatal development in Arabidopsis (Hara *et al*. 2007). An ectopic expression cassette was designed to mis-express EPF1 in sorghum, and was designated pPTN1337. The sorghum homolog of AtEPF1 (NM_127657), Sobic006G233600.1/SbiTx43006G248600.1, ORF was synthesized (GeneScript, USA), which incorporated the gene model’s 5’ and 3’ UTR elements. The synthesized ORF with its corresponding UTRs, was subcloned between the sugarcane Ubi4 promoter (Wei *et al*. 2003) and the 3’ UTR of cauliflower mosaic virus 35S transcript (T35S). The derived expression cassette was subsequently cloned into the binary vector of pPZP212 (Hajdukiewicz *et al*. 1994) and the resultant vector designated pPTN1337 (Supp. Fig. 1a).

A second vector was designed for the expression of a fusion peptide that comprises a domain from STOMAGEN (EPFL9; Hunt *et al*. 2010, Kondo *et al*. 2010) and EPF2 (Hara *et al*. 2009). This fusion consists of AA residues 26-219 from the sorghum homolog of AtEPF2 (NM_103147), Sobic006G104400.1/SbiTx43006G109800.1 that resides downstream of residues 24-37 of the sorghum homolog of STOMAGEN (AtNP_193033.1), Sobic003G299800.1/SbiTx43003G4311400.1. The fusion element is imbedded within the 5’ and 3’ UTR of sorghum EPF2 gene model (Supp. Fig. 2). This element was synthesized (GenScript, USA) and subsequently subcloned between the sugarcane Ubi4 promoter and T35S terminator. The expression cassette was then cloned into pPZP212 (Hajdukiewicz *et al*., 1994) and the derived binary vector designated pPTN1338 (Supp. Fig. 3a).

The derived binary vectors were introduced into *A. tumefaciens* strain NTL4/pTiKPSF_2_ (Luo *et al*., 2001) and the resultant transconjugants used to transform the grain sorghum genotype Tx430 as previously described (Howe *et al*., 2006, Guo *et al*., 2015)

### Initial genotyping and phenotyping of transgenic events

Progeny derived from selfed lineages of the obtained transgenic events were assessed for: (1) presence of the plant selectable marker allele via an NPTII ELISA assay according to the manufacturer’s instructions (Agdia Inc., Elkhart, IN) and (2) phenotyped for changes in stomatal density by optical tomography (Xie *et al*. 2021, see below for details). Phenotyping was performed on 15 independent, positive events of SbEPF1 and eight independent, positive transgenic events of SbEPF_syn_. Of these, two independent events carrying the SbEPF_syn_ allele, with significantly reduced stomatal density were selected for further characterisation (Figure 1a).

**Figure 1.**
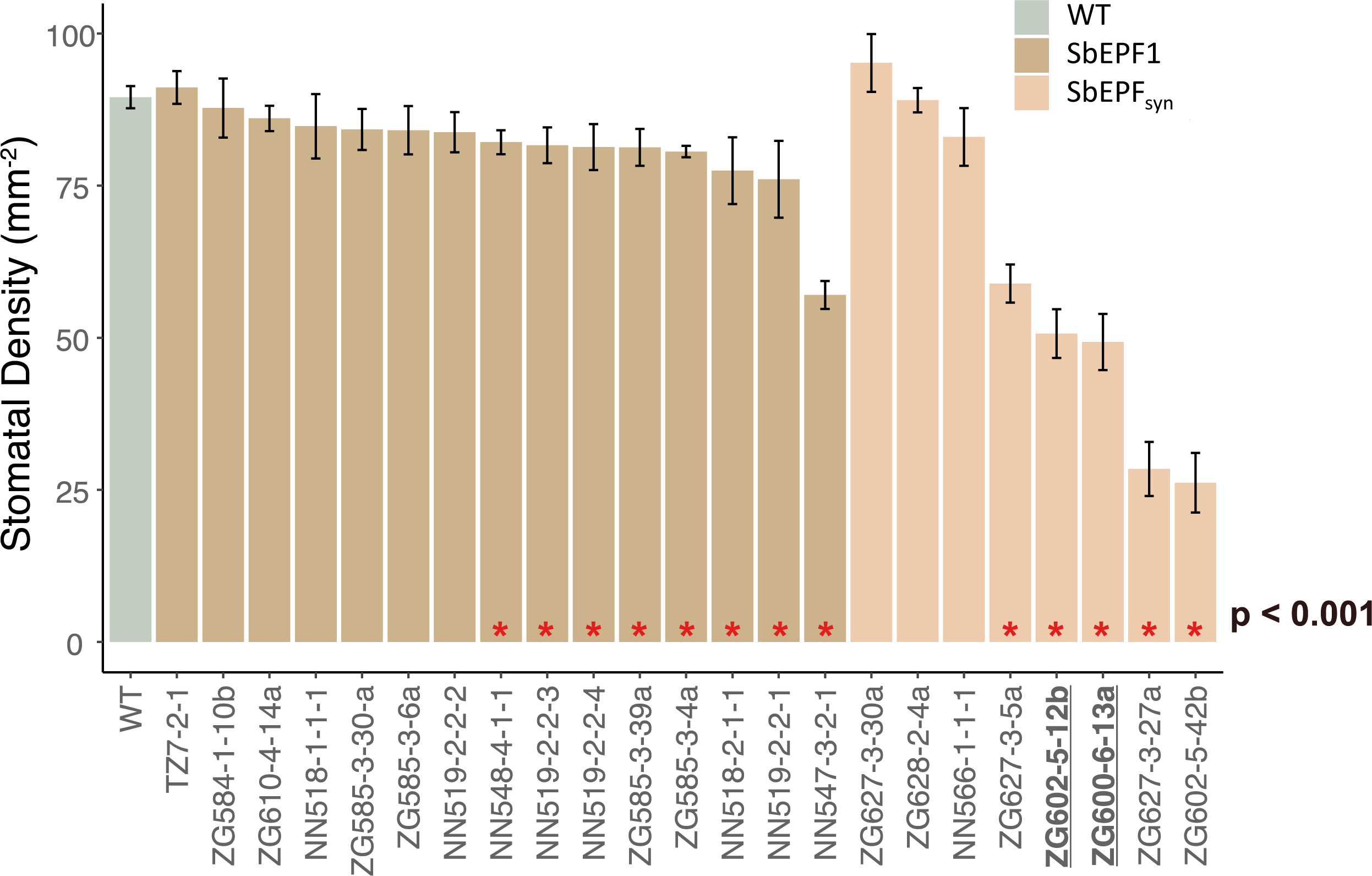
Stomatal density of independent transgenic events of **(a)** SbEPF1 and **(b)** SbEPF_syn_. The events that displayed significantly different stomatal densities compared to the WT (according to a post-hoc Tukey test following a one-way ANOVA) are denoted via red asterisks. The independent events of EPF_syn_ that were carried forward for further investigation are highlighted in bold and underlined (ZG602-5-12b and ZG600-6-13a).

The subset of the transgenic events, including the two SbEPF_syn_ events, were characterized by both Southern blot and RNA gel blot analyses as previously described (Howe *et al*. 2006, Mall *et al*. 2011). Here, total genomic DNA was digested with the *Eco*RI for Southern blot analysis (pPTN1338 events), and the membranes for both northern and Southern hybridizations were probed with a ^32^P-labeled 719 bp element that carried a region of the fusion ORF of pPTN1338 (Supp. Fig.3b,c). For the RNA gel blot analysis conducted on a set of pPTN1337 events (Supp. Fig.1b), the membrane was hybridized with an approximate 570 bp region of the SbEPF1 ORF spanning into the 3’ UTR.

### Experimental design for detailed evaluation of SbEPF_syn_

Plants were planted, grown, and phenotyped within the greenhouse facility at the University of Illinois at Urbana-Champaign (latitude 40.11°, longitude -88.21°). The greenhouse conditions were set to a 16h photoperiod (7AM-11PM) with supplementary light provided by high pressure sodium and metal halide growth lamps. The target day/night temperature was set to 28/21°C.

Wildtype (WT/Tx430) and T_2_ transgenic lineages were sown directly into trays of 4-cm deep cells filled with Sunshine™ organic germinating mix (SunGro, Agawam, MA). At the three-leaf stage, presence of the transgene was verified in the manner described previously. Additionally, T-DNA copy number analyses of NPTII was performed relative to a known single-copy sorghum gene amplicon by iDNA genetics (Norwich, UK), which confirmed the homozygosity of the transgenic allele in the respective progeny lineages going forward .

Ten replicate plants from both WT and the two independent events of SbEPF_syn_ (ZG602-5-12b and ZG602-6-13a) were transplanted into 17.5L pots containing a known mass of Sunshine Mix #4 professional growing mix (SunGro, Agawam, MA). The mass of each pot and the soil within each pot was recorded to later allow calculation of volumetric relative soil water content (% rSWC), as previously described (Ferguson *et al*. 2018). Apart from 9 days mid-growing cycle when water was withheld to perform a “dry-down” experiment (details below), plants were kept well-watered and supplemented with liquid Nature’s Source 3-1-1 NPK fertiliser (Ball DPF LLC, Sherman, TX) bi-weekly. To avoid potential spatial bias, the positions of the pots were randomised within blocks of the greenhouse space every three days across the full duration of the study, as well as every day during the “dry-down” portion of the study.

Phenotypic sample and data collection occurred in three phases. First, at the sixth leaf stage, the three most recently fully expanded leaves (i.e. leaves 4, 5, and 6) were assessed for stomatal density. At that time, light-saturated rates of leaf photosynthetic gas exchange, photosynthetic capacity, nitrogen (N) content, and specific leaf area were also measured on the youngest, fully-expanded leaf. Second, at the ninth leaf stage, a “dry-down” experiment was performed by withholding water for nine days from the plants of each genotype. Two days prior to withholding water, whole-plant leaf area was assessed. During the “dry-down” experiment, whole-plant water use and photosynthetic leaf gas exchange were measured daily. Lastly, at plant maturity, above-ground biomass production was determined.

### Leaf gas exchange measurements

When leaf six was recently fully expanded, the response of net photosynthetic CO_2_ assimilation (*A_N_*) to the concentration of leaf intercellular CO_2_ (*c_i_*) was measured using a LI-COR 6800 infrared gas exchange system equipped with a standard 6cm^2^ cuvette (LI-COR Inc., Lincoln, NE). Data collection occurred between 0800-1500, just prior to harvesting tissue from the same leaves for other leaf physiological traits. Environmental conditions were set at: 27°C, 65% relative humidity (RH), 1800 μmol m^-2^ s^-1^ photosynthetic photon flux density (PPFD), 400 µmol mol^-1^ CO_2_ concentration, and 400 µmol s^-1^ flow rate. After full photosynthetic induction was established, *A_N_*, stomatal conductance (*g_s_*), and *c_i_* were recorded as the leaf was exposed to a series of stepwise changes in sample CO_2_ concentrations of: 400, 200, 50, 150, 300, 400, 500, 600, 700, 800, and 1200 µmol mol^-1^. A custom R function was used for modelling *A_n_*-*c_i_* response curves following (von Caemmerer, 2000) to estimate the maximum rate of carboxylation by PEPC (*V_pmax_*) and the asymptote of the *A_N_*-*c_i_* curve (*V_max_*), as described previously (Markelz *et al*. 2011).

During each day of the water withdrawal experiment, *A_N_* and *g_s_* were measured everyday between 0830-1300 on the youngest fully expanded leaf. Measurements were performed using LI-6400 gas exchange systems (LI-COR Inc., Lincoln, NE) equipped with a 2cm x 3cm LED cuvette, and conditions in the gas exchange cuvette were as described for the *A_N_*-*c_i_*response measurements.

### Leaf stomatal density, SLA, N content

A Nanofocus µsurf explorer optical topometer (Nanofocus, Oberhausen, Germany) was used to assess stomatal patterning, as described previously (Haus *et al*. 2015; Ferguson *et al*. 2021). When leaf six was recently fully expanded, four fields of view (800 x 800 µm each in size) arranged in a transect between the mid-rib and margin at a position midway along the length of leaves four, five and six, were scanned at 20x magnification on both the abaxial and adaxial surfaces. 3D reconstructions were converted to 2D grey-scale images for analysis. Stomatal density was determined using the cell counter feature in ImageJ (Abràmoff *et al*. 2006) At the same time, leaf discs were sampled from the sixth leaf for estimation of specific leaf area (SLA) and tissue nitrogen (N) content, as described previously (Markelz *et al*. 2011).

### Whole-plant water use

To estimate water consumption during the “dry-down” experiment, plants were soaked to ∼100% rSWC and subsequently not watered for nine days. Each pot was weighed daily during this period. These data were used to calculate rSWC whilst accounting for pot weight at the start of the experiment, and plant mass measured on three replicates harvested on the day that water withholding started.

### Above-ground biomass production

Two-days prior to the water withdrawal period, whole plant leaf area was determined as the sum of the width multiplied by the length of every leaf. This non-destructive estimation of leaf area is highly correlated to conventional measurements of leaf area in maize (Pearce *et al*. 1975).

At full maturity, plants from both watering treatments were harvested just above the soil level and dried at 60°C for two weeks before being weighed.

### Statistical analyses

To test for overall phenotypic differences from WT in both the initial screening of all transgenic events, and the subsequent detailed evaluation of events ZG602-5- 12b and ZG602-6-13a for SbEPF2_syn_ under well-watered conditions, a one-way analysis of variance (ANOVA) comparison of means test was performed. To then determine which events were significantly different from the WT, a post-hoc Tukey test was performed.

To test for differences in genotype effects on stomatal density between leaf positions (i.e. leaf four, five or six), and to test for differences in genotype effects on above-ground biomass between watering treatments (i.e. full-watered at all times versus plants that experienced the “dry-down” treatment), two-way fully factorial ANOVA tests were performed.

To determine genotypic differences in the response of rSWC, *g_s_*, and *A_N_* to declining water availability, two-way fully factorial ANOVA tests were performed where time was treated as a repeated measure. Post-hoc Tukey tests were subsequently performed to ascertain on which days and between which genotypes significant differences occurred.

All ANOVA tests were performed using the base lm() function in R. Where multiple sub-samples were measured within a replicate plant i.e. stomatal density from multiple field of view per leaf, an average was calculated for the replicate plant and this was the input for all statistical tests. Least-squares means and standard errors for all groups from each test were computed using the lsmeans() function from the lsmeans R package (Lenth, 2016). The least-square means and standard errors are reported in the associated bar and line plots. Post-hoc Tukey tests were performed using the HSD.test() function from the agricolae R package (Mendiburu *et al*. 2015). All figures were produced using the ggplot2 R package (Wickham, 2009) with post-processing in Affinity Designer (Serif, Nottingham, UK).

## Results

### Peptide alignments of Arabidopsis and Sorghum epidermal patterning factors SbEPF1, SbEPF2, and SbEPF9

Global alignments of the sorghum gene models that encode for the epidermal patterning factors, EPF1, EPF2, and EPF9, with their corresponding Arabidopsis homologs reveal varying degrees of homology at the protein level. The sorghum and Arabidopsis EPFL9 peptides share approximately 47% identity with 66% similarity (Supp. Fig 4), with the highest degree of identity about the C-terminal region, about the core motif of the protein. An alignment of the sorghum and Arabidopsis EPF1 peptides display about 25% identity, with 31% similarity, with the highest degree of relationship also at the C-terminal region of the peptide (Supp. Fig. 5). The respective EPF2 proteins show 28% identity and 41% similarity, and like the other epidermal patterning factors, the shared alignments occur at the C-terminal region of the peptide (Supp. Fig. 6).

### Phenotypic and molecular screens of SbEPF1 and SbEPFsyn

Eight of the fifteen independent events of SbEPF1 had significantly reduced abaxial stomatal density compared to WT. However, in all but one case (NN547-3-2- 1, -26% relative to WT) the reductions in stomatal density were very modest (-8 to - 15%; Fig. 1). By contrast, five of the eight EPF_syn_ events had significantly lower stomatal density than WT, and all five had much greater reductions in stomatal density(-34 to -71%; Fig. 1). Two independent events (ZG602-5-12b and ZG600-6-13a), each carrying a single homozygous copy of the transgene and with similar reductions in stomatal density compared to WT (-43 and -45%), were selected for further investigation at the T2 stage. These lines are referred to as “12b” and “13a”.

### Stomatal patterning and light-saturated leaf gas exchange of EPF_syn_

In both lines of EPF_syn,_ stomatal density was confirmed to be significantly lower than WT on both the abaxial (p <0.001) and adaxial leaf surfaces (p < 0.001) of fully-expanded leaves at positions four, five and six on the main culm (Fig. 2). The average reduction in SD was greater on the adaxial surface (-61% in line 12b, -36% in line 13a) than on the abaxial surface (-43 % in line 12b, -30% in line 13a), and this effect was consistent across leaf positions. All genotypes had significantly greater stomatal density on the abaxial versus adaxial leaf surface.

**Figure 2.**
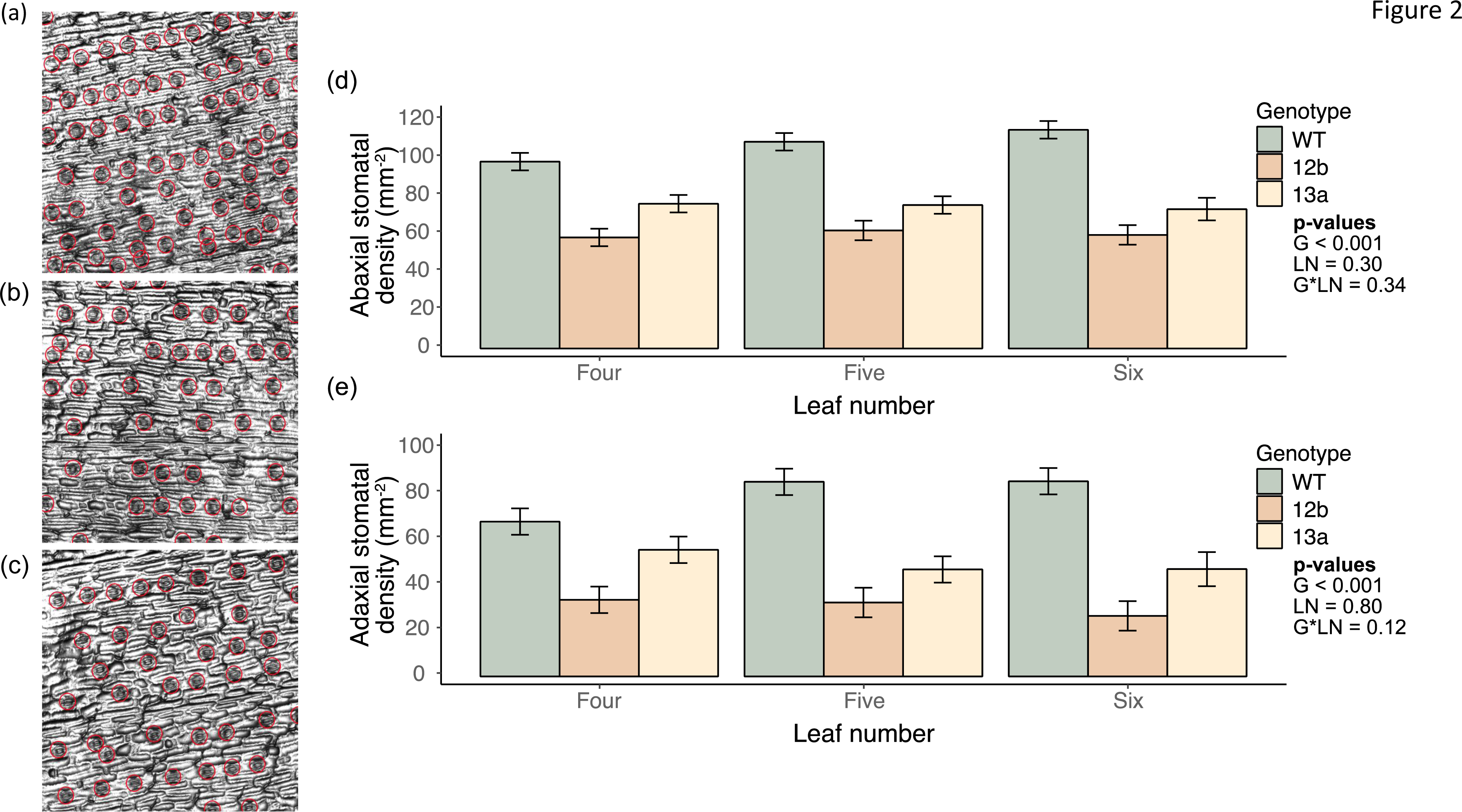
Representative micrographs of the abaxial leaf surface of: **(a)** WT, **(b)** ZG602-5-12b and **(c)** ZG600-6-13a, along with the associated **(d)** abaxial stomatal density and **(e)** adaxial stomatal density at three leaf positions on the main culm (three, four and five) for each genotype. Bars represent least square means of stomatal density for each grouping, where the errors bar represent the associated standard errors. For the abaxial and the adaxial surface, the p-values from each term in a two-way ANOVA with an interaction term are inset.

Steady-state, light-saturated g_s_ was significantly lower in EPF_syn_ compared to WT, with line 12b showing a stronger effect (-32 %, p<0.01) than line 13a (-18 %, P<0.05; Fig. 3a). In line 12b, the stronger reduction in g_s_ was accompanied by a reduction in *A_N_* compared to WT (-22 %, p<0.01; Fig. 3b). For both lines of EPF_syn_, the reduction in *g_s_*led the ratio of A_N_ to g_s_, (i.e., *iWUE*) to be significantly greater than for WT (20% for line 12b, 13% for line 13a; p<0.01; Fig. 3c).

**Figure 3.**
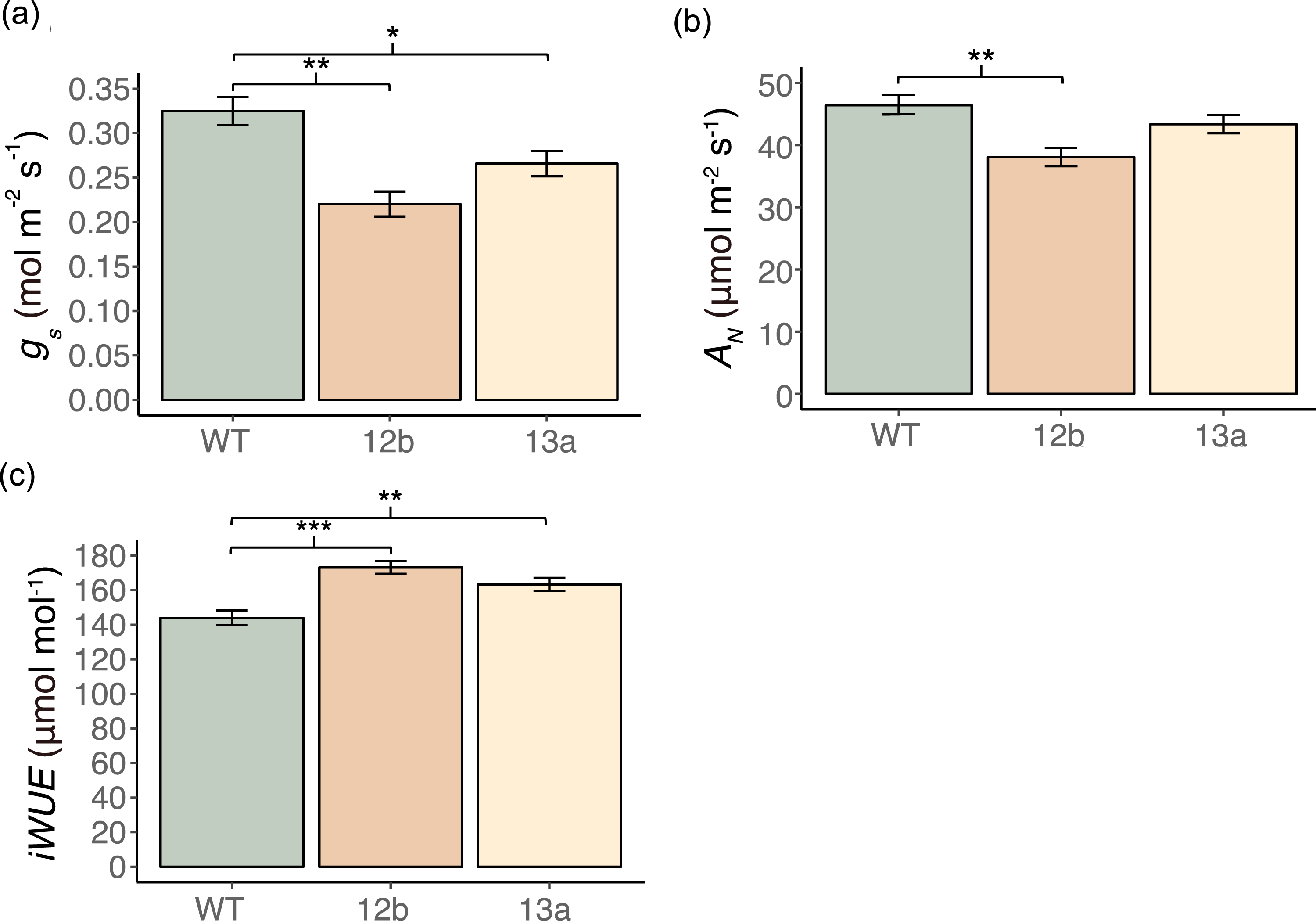
Light-saturated **(a)** stomatal conductance (*g_s_*), **(b)** net photosynthetic assimilation of CO_2_ (*A_N_*), and **(c)** Intrinsic water-use efficiency (*iWUE*) of the sixth leaf on the culm when it was the youngest fully expanded leaf of WT, ZG602-5-12b and ZG600-6-13a. Bars represent least square means and error bars represent associated standard errors. Significant differences between genotypes are denoted as * > 0.05, ** > 0.01, or *** > 0.001

The observed changes in light-saturated gas exchange of EPF_syn_ versus WT were the result of changes in both stomatal and mesophyll limitations to photosynthesis. As expected, stomatal limitation to A_N_ was low (0.04; Fig. 4a) in WT, and was unchanged in line 13a (0.04; Fig. 4c). This corresponded to the operating point of photosynthesis (blue dot) remaining above the inflection point on the A/ci curve when *g_s_* was more moderately reduced in line 13a compared to WT. In line 12b, with its stronger reduction in stomatal density, stomatal limitation to A_N_ was greater (0.08; Fig. 4b) than WT and the operating point of photosynthesis was at the inflection point on the A/ci curve. EPF_syn_ plants of both lines had lower photosynthetic capacity than WT as a result of significantly reduced apparent capacity for carboxylation by PEPC (*V_pmax_*; -32 in line 12b, -21 % in line 13a, p<0.01; Fig. 4d) and also a marginally significant reduction in the combined apparent capacity for carboxylation by Rubisco and PEP regeneration by PPDK (*V_max_*) in line 12b (-20 %, p=0.11; Fig. 4e). There were no significant differences between EPF_syn_ lines and WT in SLA (p = 0.25; Fig. 4f) or leaf N concentration (p = 0.26; Fig. 4g).

**Figure 4.**
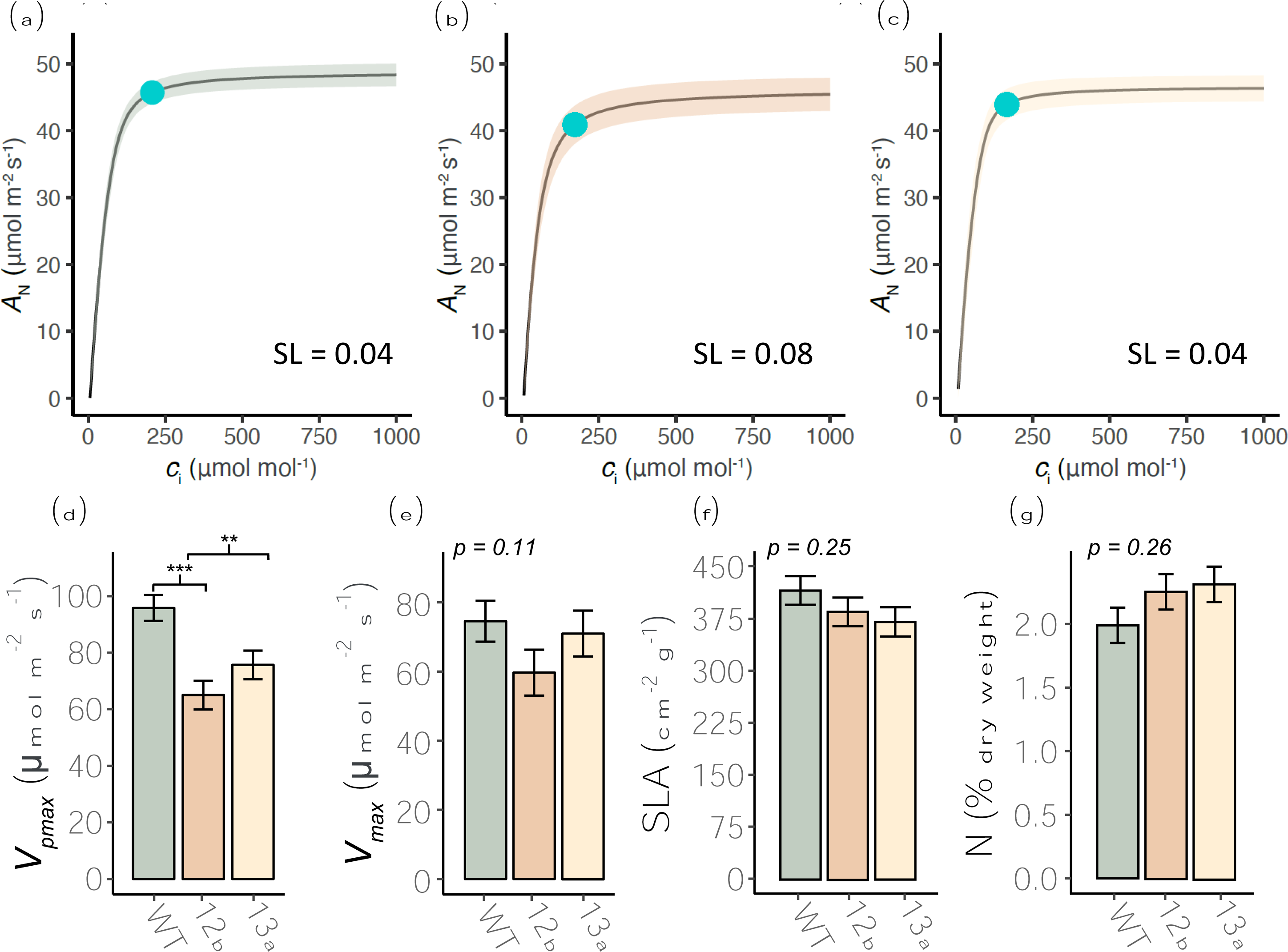
Fitted average *A-ci* response curves of **(a)** WT, **(b)** ZG602-5-12b and **(c)** ZG600-6-13a, along with **(d)** apparent maximum rate of carboxylation by PEPC (*V_pmax_*), **(e)** the asymptote of the *A_N_-c_i_ curve* (*V_max_*), **(f)** specific leaf area (SLA), and **(g)** leaf nitrogen (N) content of the sixth leaf on the culm when it was the youngest fully expanded leaf. For A/ci curves, the mean fit is represented by the solid line and the standard error is denoted by the shaded area. The stomatal limitation to A_N_ at ambient [CO_2_] (SL) is reported for each treatment. The operating point of photosynthesis at ambient [CO_2_] is shown as a blue dot on the A/ci curve. Bars represent least square means and error bars represent associated standard errors. The p-values from associated one-way ANOVA tests are inset. Where a significant effect was detected the differences between the transgenic lines and the WT according to post-hoc testing is shown.

### Interactions between leaf gas exchange, whole plant water use and drought stress

Relative soil water content (rSWC) was significantly greater in pots of both EPF_syn_ lines compared to WT as soon as day 2 of the dry-down experiment (Fig. 5a). This water saving increased in magnitude until day 6, at which point WT plants had almost exhausted the available soil water and their rate of soil drying slowed dramatically. Since initial rates of soil drying were 30-34% lower in EPF_syn_ lines compared to WT, they were sustained for longer before water supply became limiting, only displaying a slight reduction in the rate of water use on day 9, at which point pot soil moisture was almost exhausted for EPF_syn_ and WT (Fig. 5a). Considering the potential influence of plant size on rates of water use, a trend towards lower total leaf area (7-12%) in the EPF_syn_ lines compared to WT was observed at the beginning of the dry-down experiment, but the effect was not statistically significant (p = 0.60; Supp. Fig. 7).

**Figure 5.**
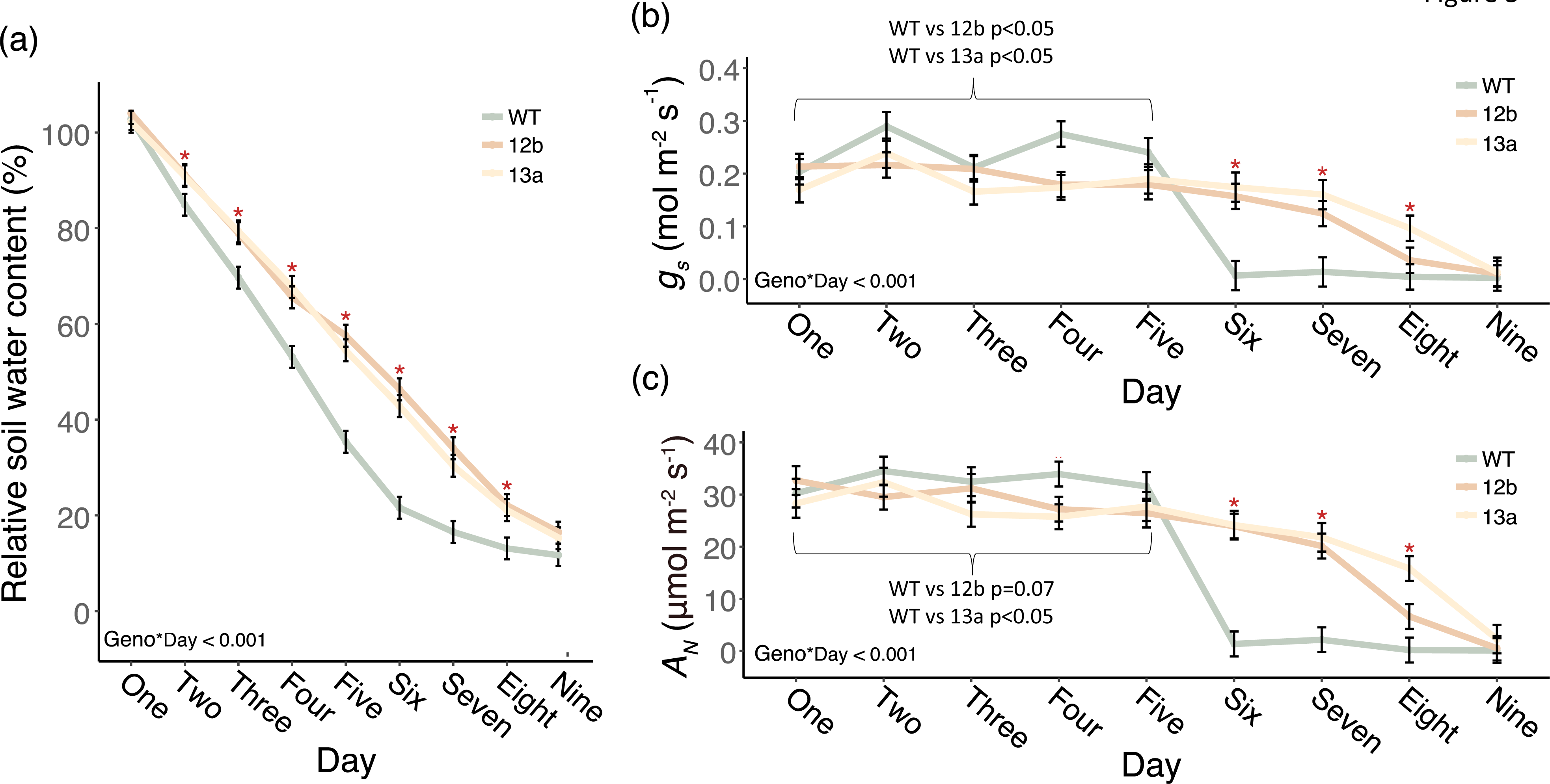
The response of **(a)** percentage relative soil water content, **(b)** stomatal conductance (*g_s_*), and **(c)** net photosynthetic assimilation of CO_2_ (A_N_) to a nine-day water withdrawal period. Points and errors bars represent least square means and standard errors, respectively. p-values associated with repeated measure ANOVAs are inset. Days where 12b and 13a were significantly different from WT for all traits are highlighted by red asterisks. The statistical results of pairwise tests between each transgenic line and the WT for a specific contrast of average *g_s_* and A_N_ on the first five days of the experiment is shown by the inset bracket.

During the first five days of the dry-down experiment, when high rates of water use were sustained in all pots, *g_s_* was 17 and 24% lower in the EPF_syn_ lines compared to WT (p<0.05; Fig. 5b). This was accompanied by more modest average reductions in A_N_ in line 12b (-10 %, p=0.07) and line 13a (-14 %, p<0.05) (Fig. 5c). On day 6, a large and rapid decline in *A_N_* and *g_s_* occurred in WT, while leaf gas exchange declined much more slowly in the EPF_syn_ lines (Fig. 5b,c). As a result, *g_s_* and AN were substantially greater for the EPF_syn_ lines compared to WT from day six through day eight (Fig. 5b,c). The difference in drought stress experienced by the plants was visually apparent with the WT being significantly wilted at the end of the “dry-down” experiment while the EPF_syn_ lines retained full turgor (Fig. 6a).

**Figure 6.**
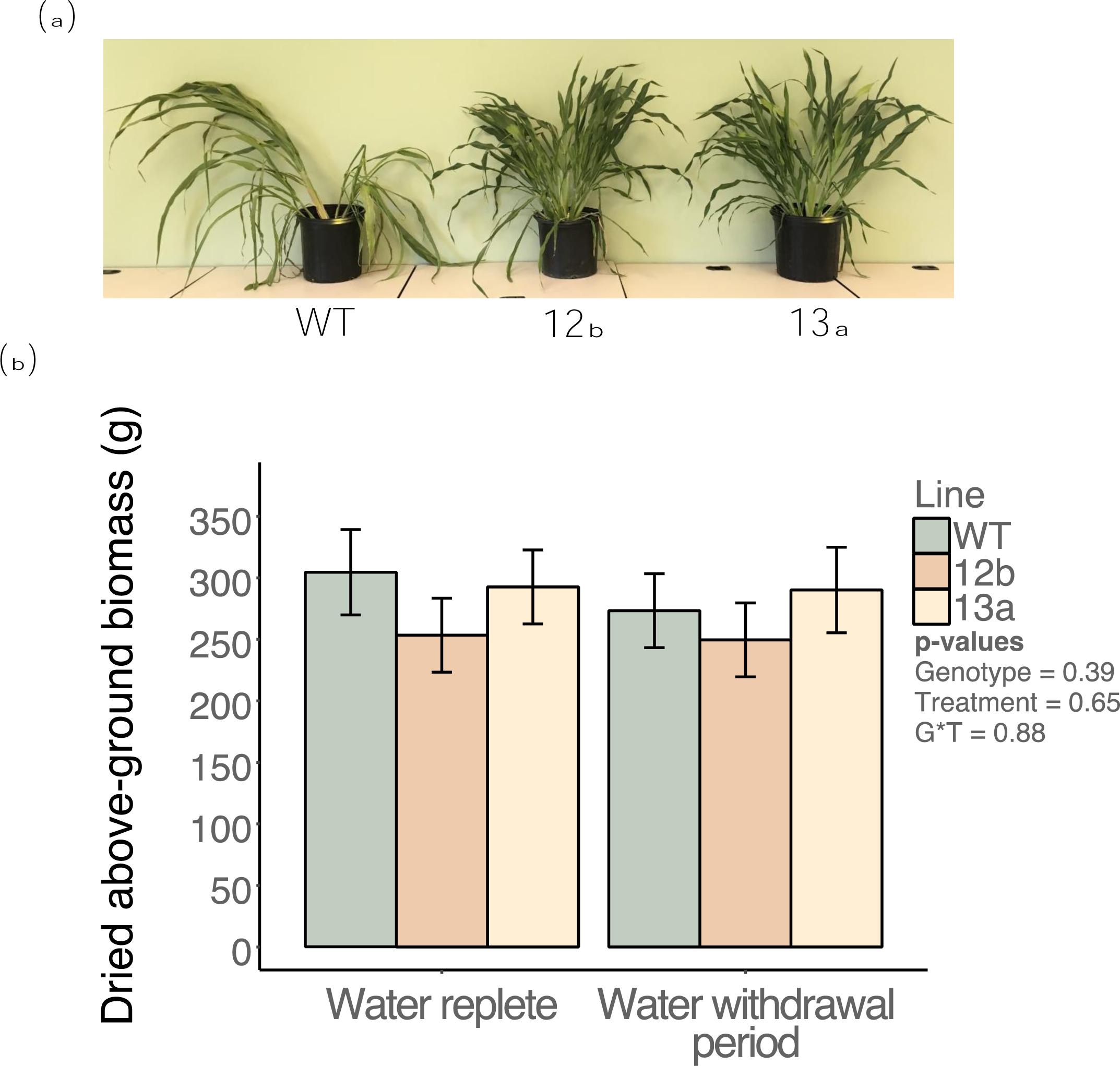
(a) Photographs of representative plants of WT, ZG602-5-12b and ZG600- 6-13a plants on day eight of the water withdrawal period. **(b)** Total dried above-ground biomass of all genotypes grown under water replete or subjected to the water withdrawal period. Bars represent least square means and error bars represent associated standard errors. p-values associated with each term from an associated two-way ANOVA are inset.

The short period of drought stress relative to the overall growing period did not significantly alter biomass production (p=0.65; Fig. 6b). Across both watering regimes, both EPF_syn_ lines had slightly shorter internode lengths and overall heights (Fig. 6a), there was no significant difference among the genotypes in total dry, above-ground biomass at maturity (p = 0.35; Figure 6b). A pleiotropic response of impaired panicle and flower development were consistently observed in the EPF_syn_ lines, resulting in significantly reduced seed production (Fig. 7). Occasionally, some leaves of EPF_syn_ lines, but not WT, would exhibit a chlorotic strip that was 2-4cm in width and contained almost no fully-developed stomata (Supp. Fig. 8).

**Figure 7.**
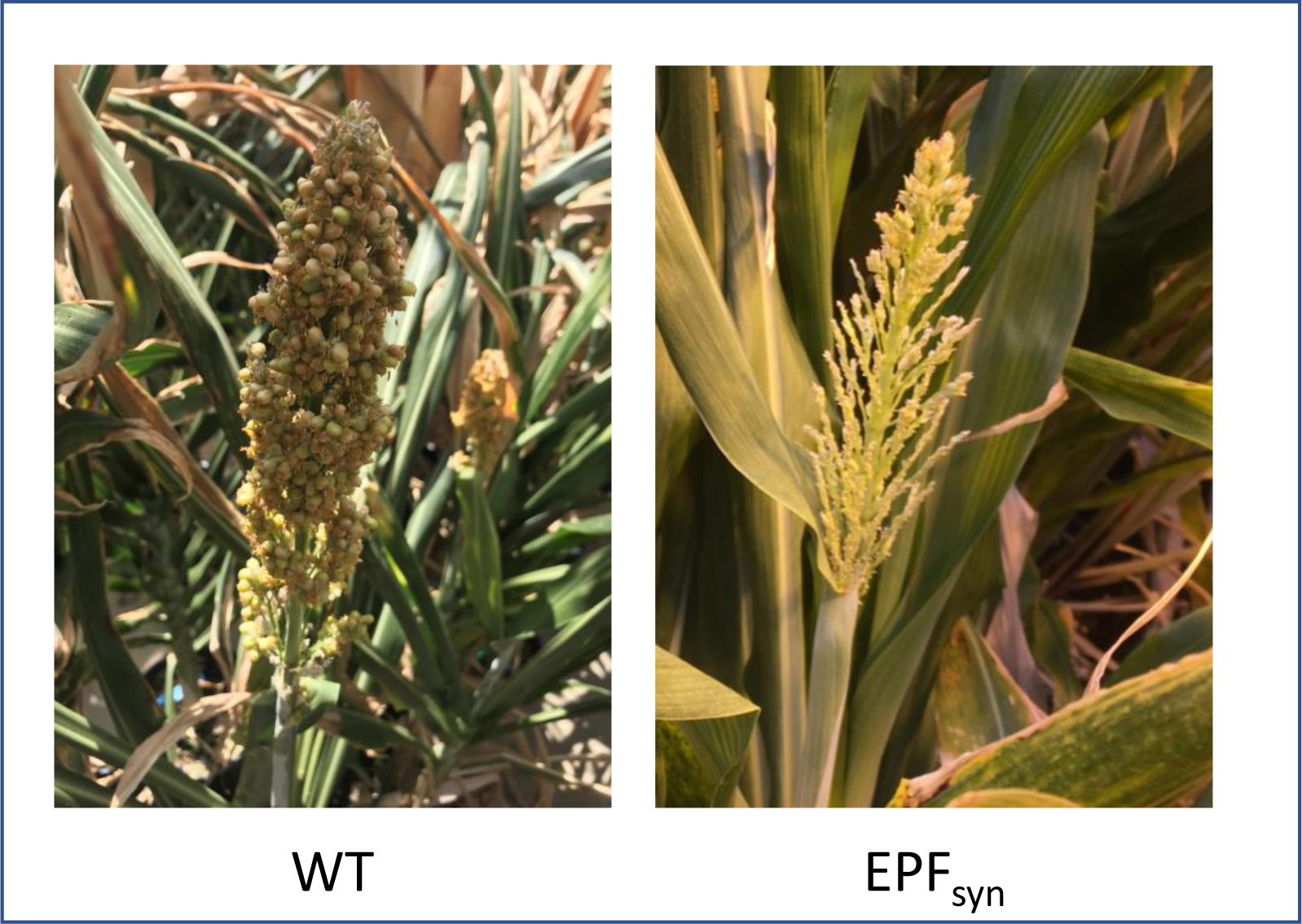
Photographs of representative panicles of WT and EPF_syn_.

## Discussion

This study contributes to our understanding of how engineered reductions in stomatal density impact leaf and whole-plant WUE in a model C_4_ species, sorghum, an important feedstock for the bioeconomy (Silva *et al*. 2022). Ectopic expression of EPF1 has been shown to significantly reduce stomatal density and increase iWUE in a number of C_3_ species (Hughes *et al*. 2017, Dunn *et al*. 2019, Mohammed *et al*. 2019). In sorghum, however, ectopic expression of EPF1 resulted in slight reductions in stomatal density (<-15%) in all but one transformation event (Fig. 1). By contrast, ubiquitous expression of EPF_syn_ resulted in reductions of -34 to -71% in stomatal density relative to WT (Fig. 1). Hence, deeper phenotypic characterizations were performed on events carrying the SbEPF2_syn_ transgenic allele to address the experimental objectives. Detailed evaluations of leaf gas exchange in two SbEPF2syn events revealed improvements in iWUE compared to WT, while highlighting the importance of achieving an intermediate reduction in stomatal density in order to lower *g_s_* while still maintaining low stomatal limitation to A_N_ (SL) (Figs. 3,4). Engineered gains in iWUE drove reductions in whole-plant water use without causing a loss in above- ground biomass production (Figs. 5,6). In addition, the study revealed the potential for pleiotropic effects on a number of developmental processes when engineering changes in stomatal development by ubiquitous expression of this SbEPF2_syn_ allele (Fig. 4,7). These observations provide a framework for future research to genetically enhance C_4_ plants in a manner that achieves the improvement in iWUE without an increase in SL, while avoiding unwanted pleiotropic effects on developmental processes other than stomatal patterning. Achieving this goal is a high priority because water-limitation is so important to global crop productivity, especially in regions where agricultural production is vulnerable to climate variability, and because improving WUE is such a valuable, if challenging, target for crop improvement (Condon *et al*. 2004; Blum 2005; Condon; Leakey *et al*. 2019).

### Balancing reductions in g_s_ with maintenance of stomatal limitation to A_N_

Analysis of A/ci curves can be used to quantify SL, which describes how much lower the observed A_N_ is compared to a theoretical A_N_ where there is no resistance for CO_2_ entry into the leaf i.e. c_i_ equals the atmospheric [CO_2_] (Farquhar and Sharkey 1982; Long and Bernacchi 2003). One of the two SbEPF_syn_ events selected for detailed characterization displayed changes in stomatal patterning and function that were close to ideal. When comparing the light-saturated leaf gas exchange of line 13a to WT, reduced stomatal density (-45%) was associated with lower *g_s_*, (-18%) and greater iWUE (+13%; Figs. 2,3). This outcome resulted first from WT sorghum having low SL (0.04; Fig. 4a), as expected for a C_4_ crop under contemporary [CO_2_] of >410 ppm (Leakey *et al*. 2019). Second, SL remained unchanged in line 13a (0.04) i.e. A_N_ was not reduced when lower stomatal density and *g_s_* limited diffusion of CO_2_ into the leaf because the photosynthetic operating point remained above the inflection point of the A/c_i_ curve (Figs. 3,4c). By contrast, SbEPF2syn event 12b had a slightly stronger phenotype, where a greater reduction in stomatal density (-69%) and *g_s_* (-32%) resulted in greater stomatal limitation to A_N_ (0.08) than in WT (0.04; Figs. 3,4b). This resulted from the operating point of photosynthesis being at a lower ci, on the inflection point of the A/ci curve, and contributed to A_N_ of event 12b being lower (-22%) than WT (Fig. 4b). These results validate the prediction, based on a leaf gas exchange model, that reducing *g_s_* to an intermediate degree can enhance iWUE with very little to no penalty to carbon (C) gain (Leakey *et al*. 2019). It is important to emphasize that the steep inflection point of the C_4_ A/c_i_ curve means that improving iWUE without a negative trade-off on photosynthetic C gain in sorghum, maize, sugarcane, miscanthus and other C_4_ grasses will depend on precisely tuning the reduction in stomatal density to achieve greater iWUE without *g_s_*and A_N_. Under the growth conditions used here, the optimal reduction in stomatal density lay somewhere between the stomatal phenotypes observed in SbEPF2syn events 12b and 13a. As future atmospheric [CO_2_] continues to rise, progressively greater reductions in stomatal density will be needed to maintain the photosynthetic operating point at the optimal location just above the asymptote of the A/ci curve.

The results of this study suggest further effort should be applied to enhance WUE in C_4_ crops by reducing stomatal conductance. While this study provides initial proof-of-concept for achieving that goal by reducing stomatal density, approaches to reduce stomatal complex size and/or the dynamic control of stomatal aperture by guard cells are also exciting possibilities for which the foundation is being laid (Des Marais *et al*. 2014, Nunes *et al*.2023). Being able to avoid an increase in SL while increasing iWUE in a C_4_ species is a distinct result from what is expected and often observed in C_3_ species (Leakey *et al*. 2019). In C_3_ crops, the lack of a photosynthetic carbon concentrating mechanism means that the A/ci curve is less steep, and photosynthesis is not expected to be CO_2_-saturated today or later this century (Leakey *et al*. 2006). Consequently, a modest loss in photosynthetic carbon gain under well-watered conditions results from engineering lower *g_s_* to achieve enhanced iWUE (e.g. Yoo *et al*. 2010, Franks *et al*. 2015, Hughes *et al*. 2017, Caine *et al*. 2019). However, crop modelling suggests this is a cost that is likely worthwhile for C_3_ crops growing in regions that are consistently water-limited (Leakey *et al*. 2019). Meanwhile, the possibility of avoiding the negative trade-off between water savings and carbon gain may allow low-*g_s_*/high-WUE C_4_ crops to enhance productivity across a wide range of growing conditions from consistently water-limited to only occasionally water-limited environments (Leakey *et al*. 2019). However, as described below, significant challenges remain to be addressed to meet that goal.

### Pleiotropic effects on mesophyll capacities for photosynthesis from ubiquitously expressing SbEPFsyn

In addition to the effects of *g_s_*, SL can be influenced by changes in photosynthetic capacity that alter the shape of the A/c_i_ curve (Markelz *et al*. 2011). Both SbEPF_syn_ events had lower apparent V_pmax_ than WT, which corresponds to a shallower initial slope on the A/c_i_ curve, and makes the inflexion point shift to greater c_i_ (Fig. 4). In event 12b, lower apparent V_pmax_ (-32 %) combined with a stronger reduction in *g_s_*to cause the increase in SL (0.08) relative to WT (0.04). In addition, there was a marginally significant (p=0.11) reduction in apparent V_max_ in event 12b, which corresponds to a lower asymptote on the A/c_i_ curve. These reductions in the mesophyll capacities for photosynthesis (apparent V_pmax_ and apparent V_max_) combined with the greater SL to explain the lower light-saturated A_N_ in event 12b compared to WT (Fig. 4).

In event 13a, the pleiotropic effects of SbEPF_syn_ on mesophyll capacities for photosynthesis were more moderate than in event 12b. Above the inflection point of the A/ci curve, where CO_2_ supply is saturating, A_N_ is limited by V_max_ i.e. a combination of Rubisco and phosphoenolpyruvate carboxylase capacity (von Caemmerer 2000). The more moderate reduction in *g_s_* compared to WT, in event 13a, meant the operating point of photosynthesis remained in this region just above the inflection point of the A/ci curve (Fig. 4c). Consequently, the lower apparent V_pmax_ (-21%) in event 13a compared to WT had no consequences for A_N_. However, a reduction in V_pmax_ is still generally undesirable from a crop performance perspective because, during drought stress, when stomata will close and c_i_ drops, photosynthesis will be operating on the initial slope of the A/ci curve (i.e. below the inflection point) and thus a lower V_pmax_ will result in reduced carbon gain. Therefore, as atmospheric [CO_2_] continues to rise, the operating point of photosynthesis will continue to shift to greater c_i_, and photosynthesis will only be operating on the initial slope of the A/ci curve during severe droughts (Leakey 2009, Markelz *et al*. 2011). So, pleiotropic effects on V_pmax,,_ while not ideal, will gradually become less of a concern in the future.

Apparent V_pmax_ is a measure of the steepness of the initial portion of the A/ci curve, which is classically defined as the apparent carboxylation capacity of PEPC (von Caemmerer 2000). PEPC catalyzes the initial carboxylation reaction of C_4_ photosynthesis in the mesophyll cells (Kanai and Edwards, 1999). The activity of PEPC is strongly correlated to leaf nitrogen content, as demonstrated recently in sorghum (Khan *et al*. 2020), where nitrogen assimilation is critical for allowing maximum photosynthesis. Moreover, in sorghum, maize, and rice it has been shown that transpiration facilitates passive nitrogen flux, with consequences for leaf nitrogen content (Niu *et al*. 2007; Matsunami *et al*. 2010; Kunrath *et al*. 2020). As such, it has been hypothesised that reducing transpiration to improve WUE might reduce nitrogen uptake and leaf nitrogen content (Hepworth *et al*. 2015), with potential consequences for the biochemical capacity of photosynthesis. However, leaf nitrogen content per unit leaf area was not significantly different between SbEPF_syn_ and WT (Figure 4g). Therefore, the observed reduction in *V_pmax_* is unlikely to be a function of this phenomena.

Alternatively, the reduced photosynthetic capacity in SbEPFsyn plants might have resulted from a signal transduction pathway triggered by expressing SbEPF_syn_ ubiquitously i.e. signaling operating in parallel to the epidermal gene network driving reduced stomatal density. In Arabidopsis, while EPF1 and EPF2 expression is focused in epidermal cells, EPFL9 is expressed in mesophyll cells and interacts with targets including light-induced factors regulating photomorphic growth (Hunt *et al*. 2010, Kondo *et al*. 2010, Wang *et al*. 2021). Leaf capacities for photosynthesis, gas conductance and hydraulic conductance need to be tightly coupled to optimize the interplay between carbon and water relations (Flexas *et al*. 2013), but the genetic underpinnings governing this physiological coordination are not well understood. The potential role of EPFs in this process would be consistent with changes in photosynthetic capacity that have been observed in some studies where expression of native stomatal patterning genes have been modified (Liu *et al*. 2015, Franks *et al*. 2015, Caine *et al*. 2019, Dunn *et al*. 2019).

In addition to the influence of PEPC carboxylation capacity, the initial slope of a C_4_ A/c_i_ curve (i.e. apparent V_pmax_) can also vary in response to changes in mesophyll conductance, which itself is determined by the resistance to CO_2_ diffusion through the internal airspaces of the leaf and then in the liquid phase from the point of dissolving in the apoplast to the point of the initial carboxylation in the mesophyll cell (Cousins *et al*. 2020). A decrease in apparent V_pmax_ could, therefore, be driven by a decrease in one or both of these components of mesophyll conductance. Such a response would be consistent with the strong positive correlation between adaxial stomatal density and mesophyll conductance observed across diverse C_4_ grass species (Pathare *et al*. 2020). Plants with a greater number of adaxial stomata per unit area demonstrated higher rates of mesophyll conductance due to an increase in the mesophyll surface area exposed to intracellular air spaces, which creates additional routes for CO_2_ diffusion to the initial site of fixation by PEPC. A pleiotropic effect of SbEPF_syn_ on mesophyll airspace development rather than photosynthetic biochemistry would be consistent with the general role of EPFs in cell fate determination and tissue development. For example, in wheat events ectopically expressing EPF1, leaf porosity was reduced alongside reductions in stomatal density and stomatal conductance (Lundgren *et al*. 2019). Further experimentation will be needed to determine if these anatomical changes are sufficient to alter mesophyll conductance, and to test if they are occurring in low-stomatal density C_4_ plants. Genes such as SCARECROW (SCR) and SHORTROOT (SHR) play roles in regulating both mesophyll and stomatal patterning in grasses (Schuler *et al*. 2017; Hughes *et al*. 2023). So, pathways that determine anatomical limitations to stomatal and mesophyll conductance can be connected. Further work will be needed to determine if mis- expression of native or synthetic EPFs can regulate both of these aspects of leaf development and function. It has been suggested that transgenes producing lower *g_s_* could be stacked with additional transgenes that confer greater mesophyll conductance, for example by making mesophyll cell walls less of a barrier to CO_2_ transport, thereby creating a positive synergistic effect on iWUE (Pathare *et al*. 2023).

### Pleiotropic effects of ubiquitously expressing SbEPFsyn on plant development

An additional and more significant off-target effect of SbEPF_syn_ was observed (Fig 7) on reproductive development, which may relate to interactions between EPF and ERECTA proteins. Bioactive EPF2 peptides are known to bind and regulate ERECTA to govern the division of protodermal cells into either pavement cells or stomatal complexes (Zoulias *et al*. 2018). ERECTA-family receptor kinases coordinate stem cell functions between the epidermal and internal layers of the shoot apical meristem, and they are demonstrated to regulate floral patterning, fertility, and organ identity in addition to stomatal patterning Arabidopsis (Shpak *et al*. 2003; Cai *et al*. 2017; Kimura *et al*. 2018). So, it is possible that the ubiquitous expression of SbEPFsyn may have perturbed equivalent pathways in sorghum, leading to impaired panicle and flower development (Fig. 7). If so, enhancing iWUE in sorghum while avoiding pleiotropic effects on photosynthetic capacity and seed production might be achieved by the use of tissue specific promoters that can limit expression of EPFsyn to the epidermis during early phases of leaf development.

### Whole-plant biomass production, water use, and drought avoidance

The overall biomass production of SbEPFsyn plants was equivalent to that of WT (Fig. 6). As described above, this appears to have been due to the pleiotropic effects of expressing SbEPFsyn being focussed on developmental processes, and under conditions of ample water supply, the anatomical consequences had mild (event 12b) to no effect (event13a) on photosynthetic carbon gain.

Total plant water use is a function of both the rate of water use per unit leaf area and the total leaf area of the plant. The rate of water use by the two SbEPFsyn events was -30 and -34% lower than WT (Fig. 5a). Most of the water savings can be attributed to the reduction in g_s_, which averaged -20 and -23% lower in the two SbEPF_syn_ events compared to WT, when averaged across all the dates of measurement on which water supply was not limiting (Fig. 5b). However, it seems likely that the -5 and -10% changes in total plant leaf area of the SbEPFsyn events compared to WT also contributed to lower rates of water use, even if they were not resolved as statistically significant (Supp. Fig. 7).

When water supply was withheld in the dry-down experiment, differences in rates of water use meant that SbEPFsyn plants took nine days to exhaust the water supply in their pots versus six days for WT (Fig. 5a). When the water supply ran out for WT plants it triggered a substantial and rapid drop in *g_s_* and A_N_ (Fig. 5b,c). It is noteworthy that *g_s_*and A_N_ declined gradually between days six to nine in the SbEPFsyn events, compared to very abruptly decrease between days five and six in the WT (Fig. 5b,c). This is consistent with the interpretation of the A/c_i_ curve data described above i.e. under mild, initial drought stress the photosynthetic operating point of SbEPF_syn_ plants required less of a drop in *g_s_* and c_i_ to sit at or below the inflection point of the A/c_i_ curve, leading to a modest loss of C gain. Nevertheless, these experimental results support the notion that a low stomatal density, low *g_s_*, high iWUE strategy in a C_4_ crop results in greater carbon gain over the dry-down period as a whole, which may enhance photosynthetic carbon gain and biomass production in locations where water supply limits productivity (Leakey *et al*. 2019). It is important to note that drought stress generally develops more rapidly and severely in pot experiments than under field conditions. For example, dry-down of the large soil volume at field locations in the Central U.S. can take weeks rather than a few days to develop (Leakey *et al*. 2004; Markelz *et al*. 2011; sorghum drought). In addition, the dynamics of dry-down and re-wetting cause plant drought stress events to vary with soil type and climatic conditions. So, field trials and crop modelling will be needed to quantify the optimal phenotype of weak versus strong reductions in stomatal density and *g_s_* across a range of growing conditions.

### Conservation of EPF function across C_3_ and C_4_ lineages

In Arabidopsis, overexpressing the native EPF1 reduces stomatal density by an average of 53% across >10 independent events (Hara *et al*. 2007). Overexpressing, the barley EPF1 ortholog in Arabidopsis also produces a substantial reduction (42% in barley across two events) in stomatal density and overexpressing the rice EPF1 ortholog in the *epf2* Arabidopsis mutant restores WT stomatal density (Hughes *et al*. 2017; Caine *et al*. 2019). This highlights the conserved functionality of EPF1 across the dicot and C_3_ monocot functional types. Accordingly, overexpression of the native EPF1 genes in barley, rice, and wheat achieves significant reductions in stomatal densities that are like those or greater than those achieved via AtEPF1 overexpression in Arabidopsis, i.e., 52% average reduction in barley across two events, 45% average reduction in rice across three events, and 70% average reduction in wheat across two events (Hughes *et al*. 2017; Caine *et al*. 2019; Dunn *et al*. 2019). The minor impact on stomatal density in response to the overexpression of SbEPF1 in this present study (Fig. 1a), compared to what has been observed in barley, rice, and wheat, is consistent with the possibility of functional divergence or redundancy of EPF1 in sorghum and possibly other C_4_ grasses. Essential developmental processes are often maintained through functional redundancy, where two or more genes essentially perform the same function, such that manipulating the expression of one of those genes has little or no effect on the prevailing phenotype (Nowak *et al*. 1997). Gene and genome duplication events can enhance the likelihood of gene functional redundancy as paralogous genes that overlap in functionality are generated (Lee *et al*. 2014). Moreover, non-homologous genes can acquire similar functions as species and clades diverge (Kafri *et al*. 2009). The hallmark Kranz anatomy in C_4_ grasses is distinct from the leaf structure of C_3_ grasses, highlighting the possibility for the evolution of functional redundancy and/or novel function acquisition across these lineages. Indeed, the recent study of (Hughes *et al*. 2019) demonstrates that Kranz cell patterning in maize is regulated in part by the redundant copies of the SCARECROW 1 (SCR1) gene that plays different roles in the root and shoot. Moreover, phylogenetic divergence in stomatal characteristics between C_3_ and C_4_ grasses has been observed to mirror the evolution of the C_4_ photosynthetic pathway and local adaption (Taylor *et al*. 2012; Lundgren *et al*. 2014). Our results regarding SbEPF1 highlight the necessity of elucidating the genetic networks that underpin stomatal development in C_4_ grasses to understand how they differ between C_3_ dicots and grasses. The focus in the current experiments on understanding the downstream effects on carbon and water relations of reduced stomatal density mean that understanding the molecular mechanism by which EPF_syn_ operated is beyond the scope of the current study. This is just one of many knowledge gaps remaining about the genetic basis for stomatal development in C4 grasses that need to be addressed.

## Conclusion

In summary, this study provides support for prediction from modelling studies that engineering to reduce stomatal conductance can improve iWUE and act to lower plant water use without reducing biomass accumulation of a model C4 crop. However, it also highlights: (1) the potential for pleiotropic effects on a range of developmental when a mobile signaling peptide is expressed ubiquitously; and (2) the currently limited understanding of the genetic basis for stomatal development in C_4_ grasses. These knowledge gaps will need to be addressed and more sophisticated engineering strategies and/or additional genes targeted in order to retain the desirable stomatal phenotypes observed here without the negative consequences of pleiotropic effects on the agronomics of the crop.

## Acknowledgements

We thank Mike Masters for assistance with elemental analysis. We thank the UIUC Plant Biology and Crop Sciences greenhouse staff for support. The information, data, or work presented herein was funded in part by the Advanced Research Projects Agency-Energy (ARPA-E), U.S. Department of Energy, under Award Number DE-DE- AR0000661. No conflicts of interest are declared.

## Supplemental material

**Supplemental Figure 1.**
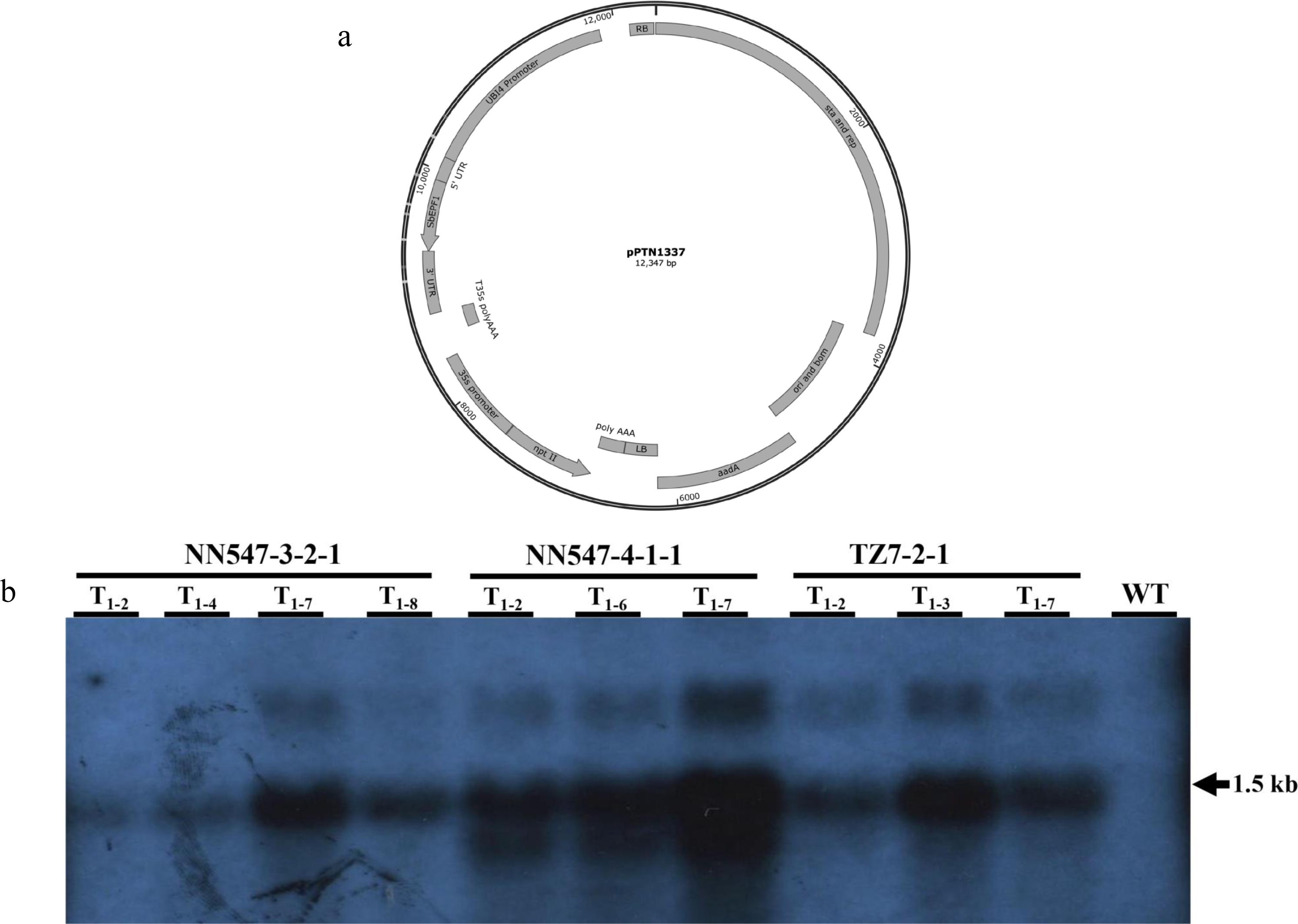
**(a)** The binary vector pPTN1337 carrying the ectopic expression cassette of the SbEPF1. Ubiquitin, UBI4, constitutive promoter from sugarcane. The 5’ & 3’ UTR elements of the gene model Sobic006G233600 delineate the SbEPF1open reading frame. The T35s polyAAA refers to the terminator of transcription from cauliflower mosaic virus. LB and RB refer to the left and right T-DNA borders. While the aadA, ori/bom, and sta/rep indicate the positions of the backbone of the broad host range binary vector that include the bacterial selectable marker (aadA), for spectinomycin resistance, ori/bom origin of replication, and basis for mobility, and stability and second origin of replication motif (sta and rep). The plant selectable marker cassette resides proximal to the LB element. This cassette harbors the cauliflower mosaic virus 35s promoter, neomycin phosphotranferase II gene from E. coli (npt II) and is terminated by the T35s polyAAA. **(b)** Northern blot analysis on sorghum events (T1) carrying the SbEPF1 expression cassette (pPTN1337). Total RNA gel was hybridized with a 570 bp element that contained a downstream region of the SbEPF1 ORF, including some of the 3’ UTR. The expected 1.5 kb signal is indicated, the observed larger hybridization signal may be associated with the unprocessed transcript that still harbors the upstream intron associated with the UBI4 promoter element. No signal was observed in the control Tx430 (WT: lane 11), likely to the relative low expression of SbEPF1, which is below detection with the hybridization assay. Lanes 1-4: T1 individuals derived from event NN547-3-2-1. Lanes 5-7: T1 individuals derived from event NN547-4-1-1. Lanes 8-10: T1 individuals derived from event TZ7-2-1.

**Supplemental Figure 2.**
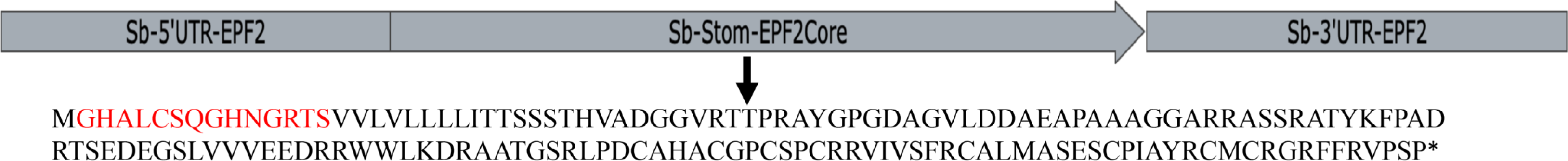
Diagrammatic representation of the translational product of the fusion peptide expression cassette of SbEPFL9/SbEPF2 present in the binary vector pPTN1138. The 14 amino acid residues shown in red at the N-terminal region of the fusion peptide from Sb stomagen (SbEPFL9) and the C-terminal amino acid residues 26-219 from the SbEPF2 in black. The respective UTR’s from Sobic006G104400.1 are represented as Sb-5’UTR-EPF2 and Sb-3’UTR-EPF2.

**Supplemental Figure 3.**
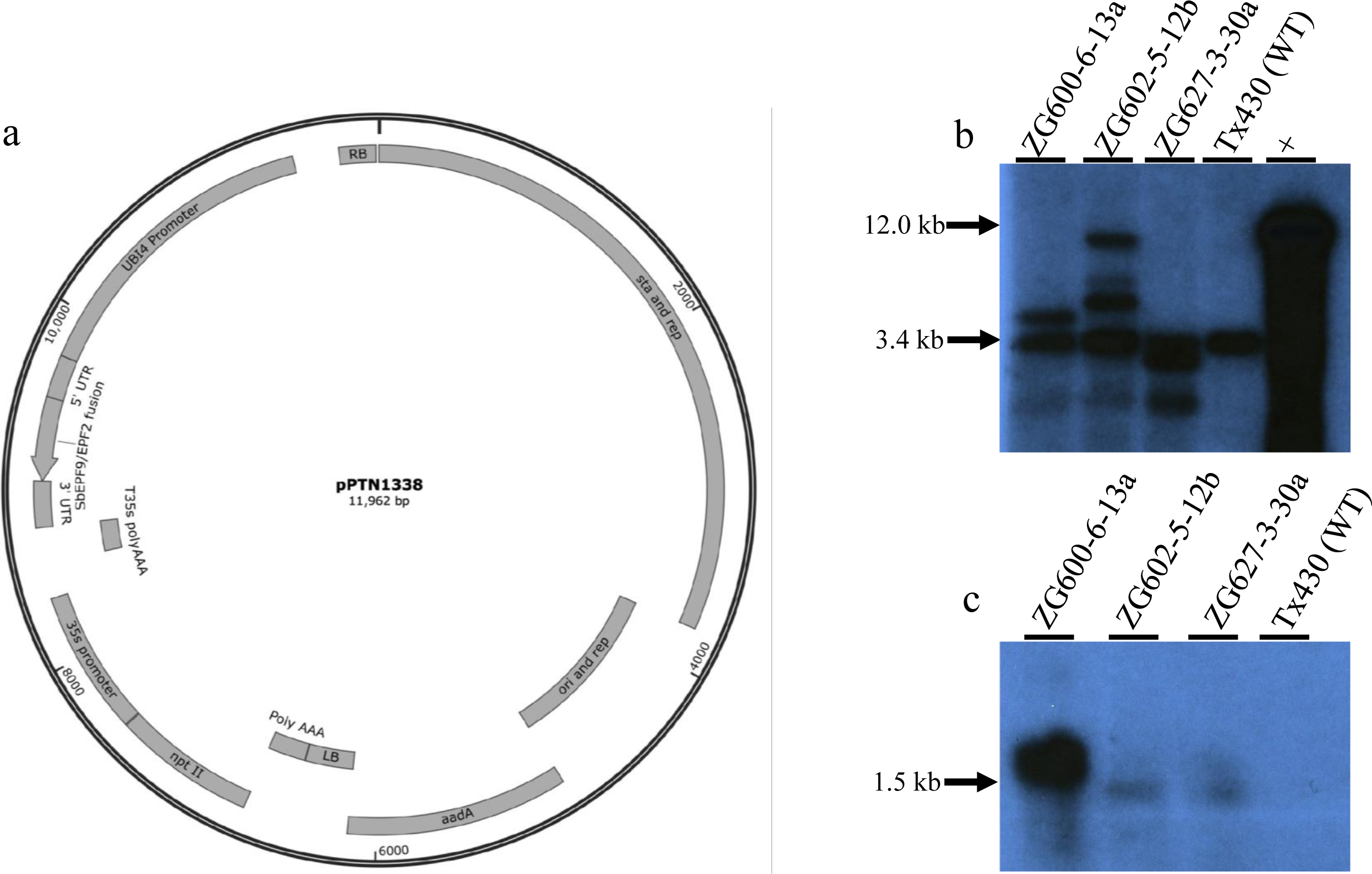
**(a)** The binary vector pPTN1338 fusion peptide expression cassette of SbEPLF9/SbEPF2, delineated by the 5’ & 3’ UTR from gene model Sobic006G104400.1 and terminated by the T35s polyAAA. Other details of design match that of pPTN1337 and described in Supp. Fig. 1. **(b)** Southern blot and **(c)** northern blot analyses on sorghum events ZG600-6-13a, ZG6002-5-12b and ZG627- 3-30a of SbEPF_syn_. (a): Southern blot restriction (EcoR1) digested total genomic DNA was hybridized with 719 bp element of the fusion ORF. The endogenous signal was observed in the control (WT) lane 4, at approximately 3.4 kb, while the events, lanes 1-3, displayed varying two to four hybridizing loci demonstrating each event is an independent event. Lane + is approximately 10 pg of vector pPtN1138 digested with EcoR1. (b): northern blot analysis on total RNA from sorghum events ZG600-6-13a, ZG6002-5-12b and ZG627-3-30a (lanes 1-3) and control (WT) lane 4. Membrane was hybridized with same probe used in the Southern.

**Supplemental Figure 4.**
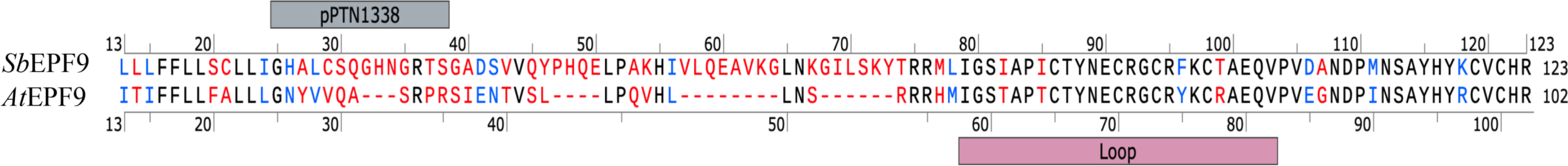
The global alignment of SbEPFL9, top strand, and AtEPFL9, bottom strand, reveal a 47% identity (black), and 66% similarity (blue). The loop motif is spanning residues 57-83, while the N-terminal fusion residues incorporated into the ORF of expression cassette that resides in the binary vector pPTN1338 is shown above residues 25-38.

**Supplemental Figure 5.**
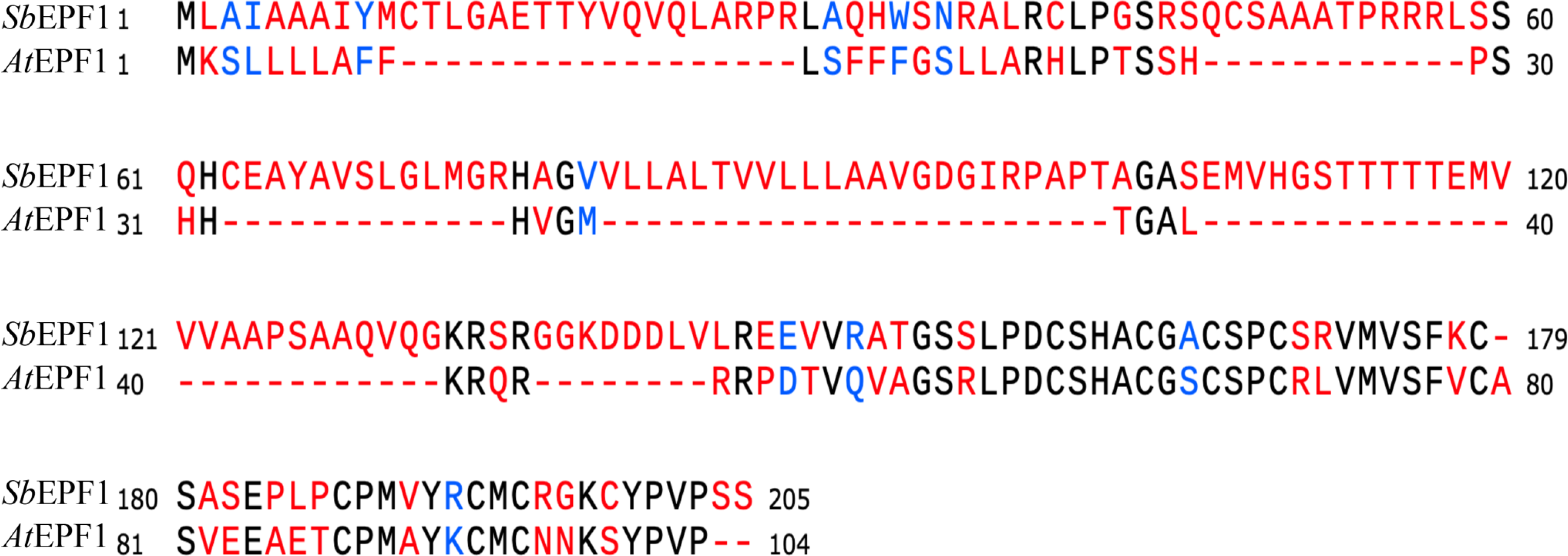
The global alignment of SbEPF1, top strand, and AtEPF1, bottom strand, reveal a 28% identity (black), and 41% similarity (blue).

**Supplemental Figure 6.**
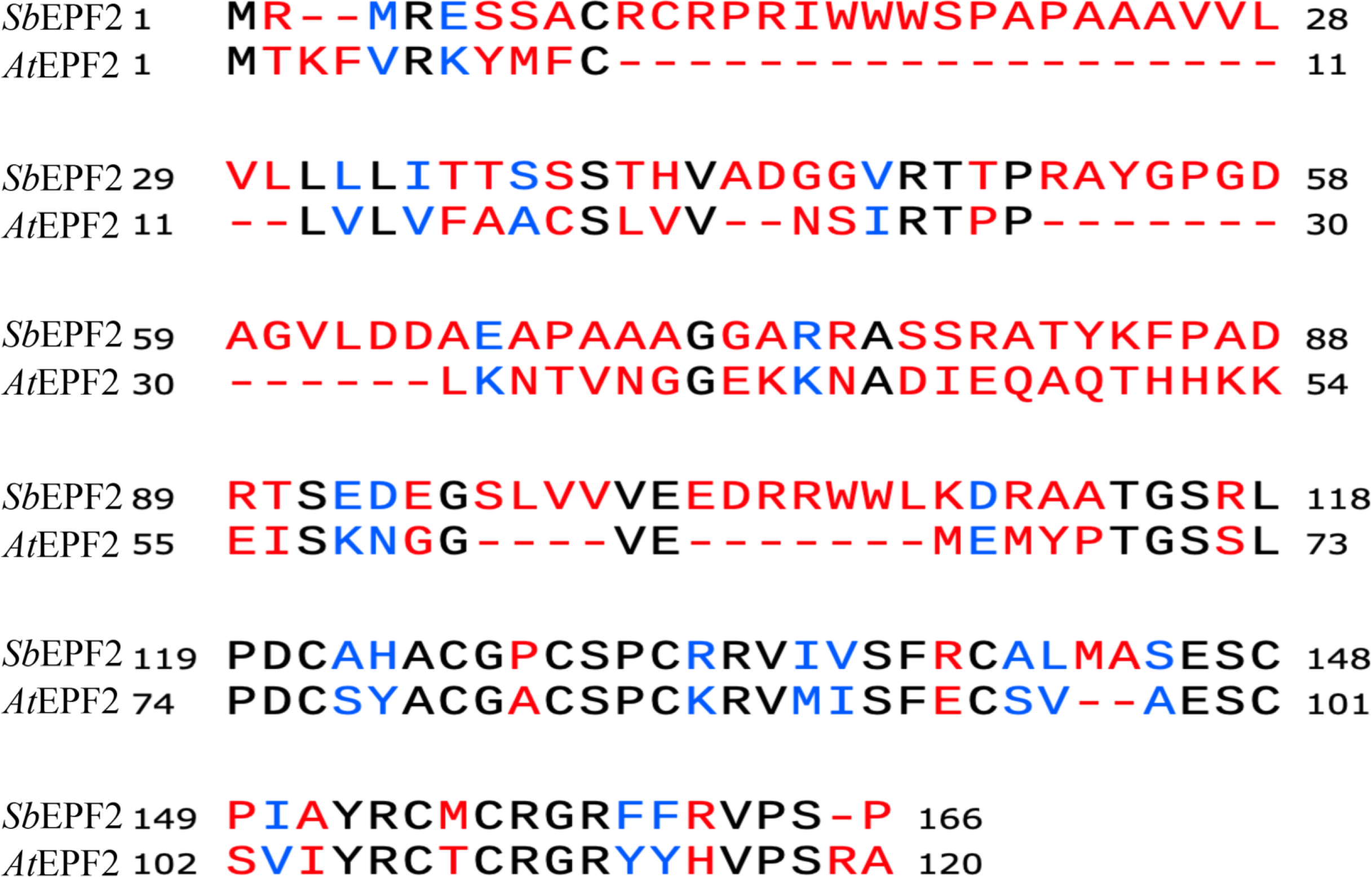
The global alignment of SbEPF2, top strand, and AtEPF2, bottom strand, reveal a 25% identity (black), and 31% similarity (blue).

**Supplemental Figure 7.**
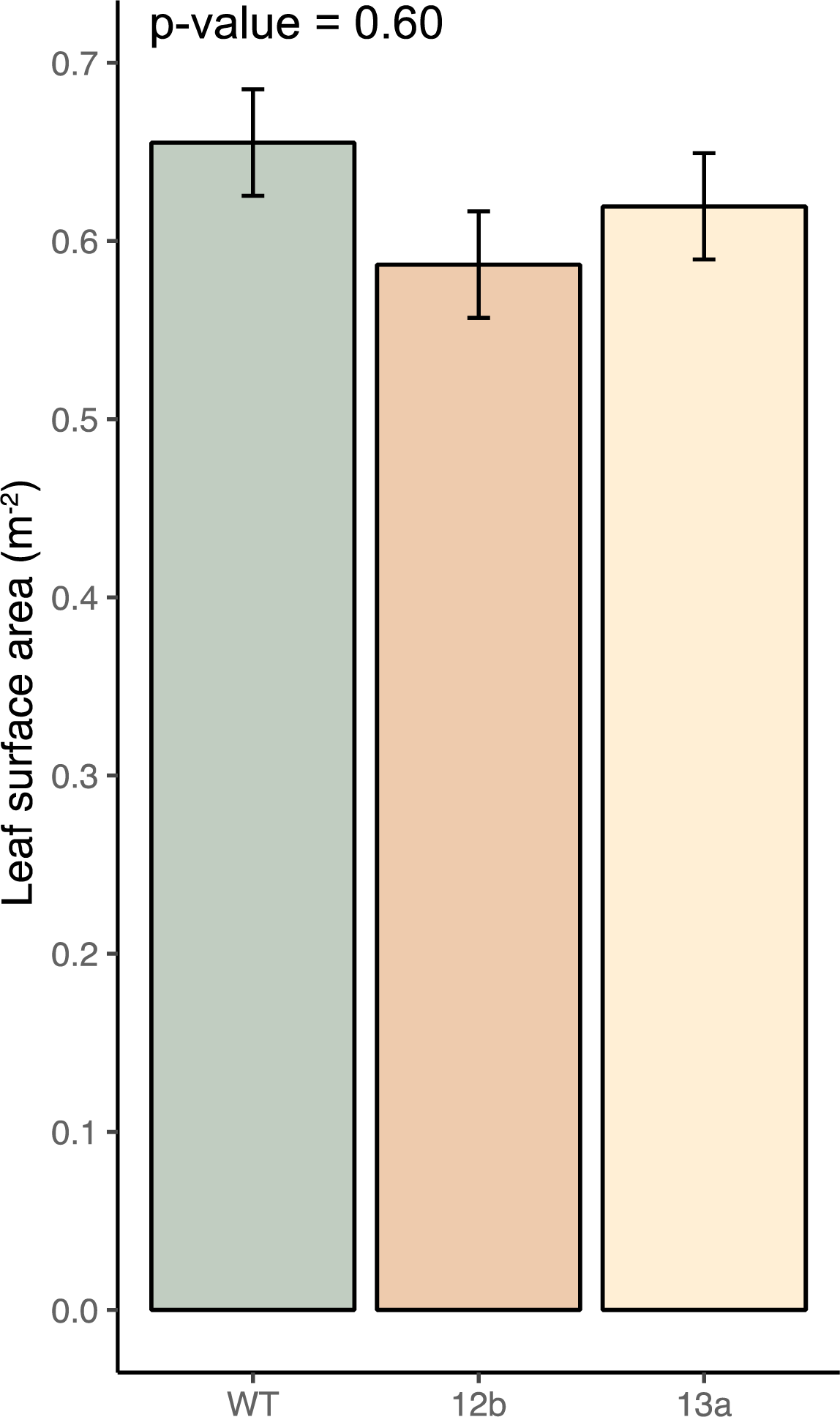
Total leaf surface area of wildtype, ZG602-5-12b and ZG600-6-13a. Bars represent least square means and error bars represent associated standard errors. P-value from ANOVA testing the effect of genotype is inset.

**Supplemental Figure 8.**
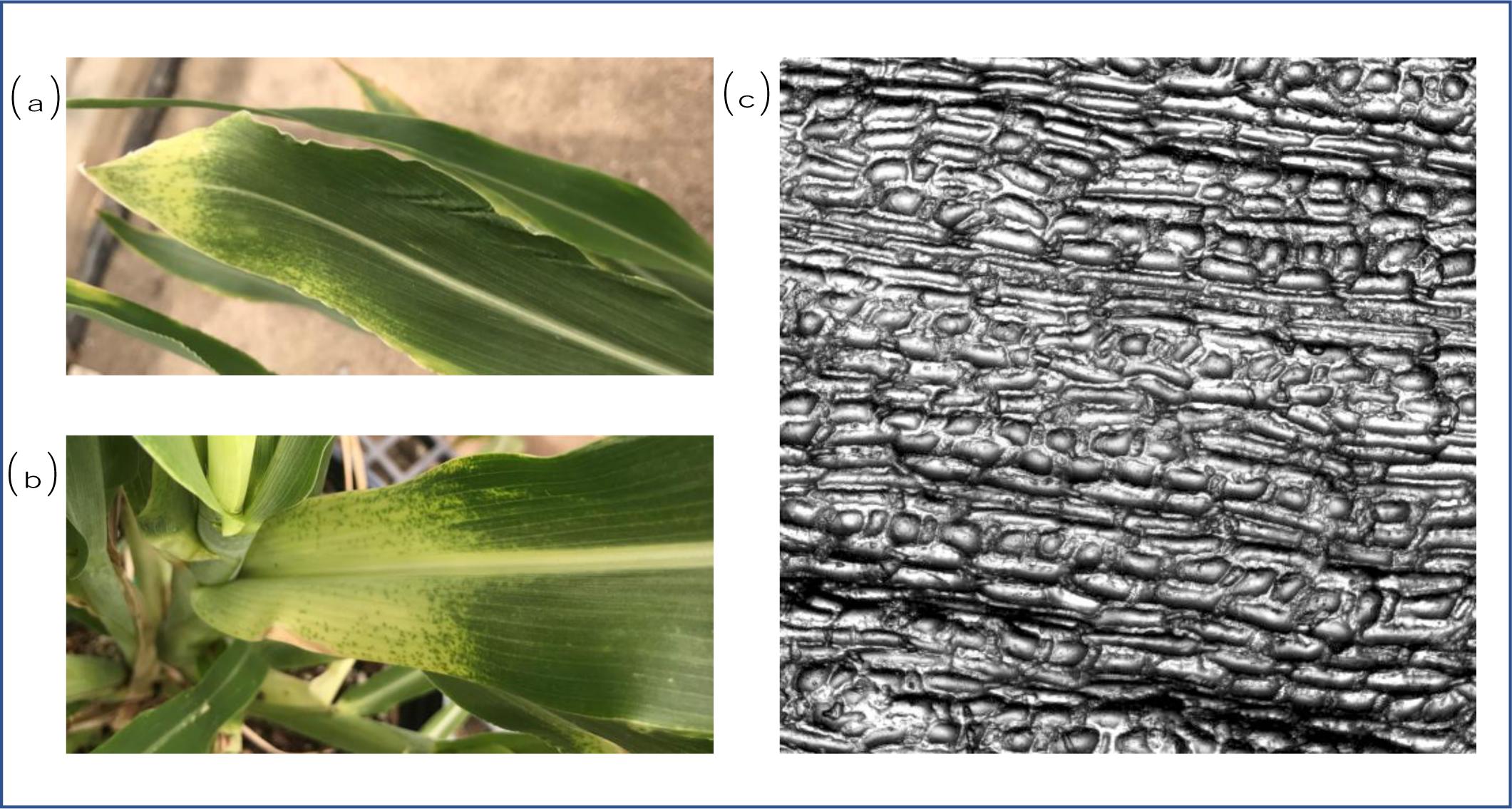
**(a,b)** Photographs of representative plants featuring the occasionally observed phenotype of a short white band on a mature leaf, which **(c)** a representative optical tomography image shows to be a region almost entirely lacking stomata on the leaf epidermis.

